# The Balancing Act: Olive baboon *(Papio anubis)* occupancy is associated with resource-related environmental variables rather than relative abundance of predators

**DOI:** 10.64898/2026.04.13.717486

**Authors:** N. van Rooyen, F. Prugnolle, V. Rougeron, T.R. Hofmeester

## Abstract

Understanding how the fear of predation acts as a driver of spatial distribution is fundamental to animal behaviour research, yet this relationship is not wholly understood in primates such as baboons. Olive baboons (*Papio anubis*) have evolved a diverse range of antipredator strategies that reduce, but do not eliminate, predation risk from the large carnivores they encounter across their broad geographic range. This raises a critical question: does the need to access essential resources outweigh the risk of predation when determining habitat selection? We addressed this question by examining the relative influence of three environmental factors and relative predator abundance on olive baboon occupancy patterns and detection probability in Serengeti National Park, Tanzania. Using data from 225 camera traps deployed by the Snapshot Safari program, we fitted three separate Bayesian occupancy models, each incorporating the same three environmental covariates (terrain ruggedness index, distance to nearest river, and Normalized Difference Vegetation Index, NDVI), together with the relative abundance of one of three principal predators (lion, leopard, or spotted hyena). This approach allowed us to assess whether environmental covariates associated with baboon occupancy remained consistent across different predator contexts. Baboon occupancy strongly increased with terrain ruggedness in all three models and consistently decreased with a greater distance to rivers. Vegetation greenness (NDVI) showed a positive association with baboon occupancy, though credible intervals narrowly overlapped zero. NDVI also showed a strong positive relationship with baboon detection probability. Associations between predator relative abundance and baboon occupancy varied between models: the relative abundance of lions and spotted hyenas showed no strong association with baboon occupancy, whereas the relative abundance of leopards was strongly correlated with baboon occupancy, consistent with shared habitat preferences. Our findings demonstrate that, independent of predator presence, olive baboon spatial distribution in the Serengeti is primarily and consistently associated with resource-related environmental features. This study expands our knowledge on the ecological factors that influence primate occupancy by showing that, for a behaviourally flexible species with diverse antipredator strategies, access to essential resources can outweigh spatial avoidance of predators in a multi-predator landscape.

## Introduction

The fear of predation is an essential component for understanding animal behaviour as fear acts as a measurable driver influencing species’ space use across a landscape (Laundré et al., 2001, 2010). Predation risk varies heterogeneously across landscapes because of differences in terrain and predator abundance, creating a mosaic of high- and low-risk zones that prey must navigate to survive (Longland and Price, 1991; Kauffman et al., 2007). In this context, the “landscape of fear” concept highlights that spatial decisions are the outcome of trade-offs between food acquisition and safety from predators (Riginos et al., 2015; Northfield et al., 2017; Gaynor et al., 2019).

As prey species are constrained by the distribution of available resources (Sih, 2005) and must therefore balance the risk of exposure with the need for food and water, predators, in turn, tend to concentrate their hunting effort where prey congregate and where landscape features provide advantages to their mode of hunting (Northfield et al., 2017). However, responses to predation risk can differ even among ecologically similar species. For instance, vervet monkeys (*Chlorocebus pygerythrus*) and samango monkeys (*Cercopithecus albogularis*) co-occurring at the same study site respond differently to identical predator species. Vervet monkey ranging behavior was significantly influenced by the perceived predation risk from chacma baboons (*Papio ursinus*) and leopards (*Panthera pardus*), but not by eagles or snakes, with food availability acting as a significant predictor of space use (Willems & Hill, 2009). In contrast, samango monkeys at the same study site were significantly influenced by aerial predators such as crowned eagles (*Stephanoaetus coronatus*) and Verreaux’s eagles (*Aquila verreauxii*), with food availability having little effect on space use (Coleman & Hill, 2021).

Due to the broad geographic range of primates across Africa (Zinner et al., 2011), different species such as olive baboons (P*apio anubis*) — a semi-arboreal (Hammond et al., 2025), highly social (Schreier & Swedell, 2012), and behaviourally flexible and aggressive species (Cowlishaw, 1994) — have evolved a diverse range of antipredator strategies in order to persist across multi predator landscapes. These strategies include vertical escape, collective vigilance, and predator mobbing (Karpanty & Wright, 2007; Cowlishaw, 1997a; Gursky, 2007; Bezerra et al., 2008; Crofoot, 2013; Bidner, 2018). While these responses reduce predation risk, they do not eliminate the threat of predation entirely. This raises a critical question: does the need to access essential resources outweigh the risk of predation when determining habitat selection in olive baboons?

Movement patterns in baboons appear to be primarily driven by seasonal rainfall, access to water sources and resource dynamics (Barton et al., 1993; Markham et al., 2015; Hammond et al., 2025), with greater ground movement occurring during dry periods. This pattern suggests that, at broader temporal and spatial scales, the acquisition of essential resources may take precedence over predator avoidance (Fehlmann et al., 2017; Suire et al., 2021; Paietta et al., 2022), with anti-predator behaviours being deployed mainly in high-risk contexts (Cowlishaw, 1994; Suire et al., 2023). However, evidence from earlier studies offers a more nuanced view. Baboons have been described as spending more time foraging in low-risk, food-poor habitats than in high-risk, food-rich ones (Cowlishaw, 1997b). Together, these findings suggest that the relative importance of food and fear in determining baboon occupancy (the proportion of an area occupied by a species, MacKenzie et al., 2005) and detection probability (the likelihood of detecting a species, if the species is present, Niedballa et al., 2015), may be context and study site dependent.

Camera trap occupancy modelling offers an alternative method to investigate this question, as camera traps are a non-invasive, continuous monitoring approach (Newey et al., 2015) that collect large scale, less detailed data simultaneously across a single or multiple study sites (Swanson et al., 2015; Moore et al., 2020; Bersacola et al., 2022). Against this backdrop, the olive baboon provides an interesting model for testing whether space use is chiefly structured by essential environmental variables or by the threat of predation from their primary predators, lions, leopards, and spotted hyenas (*Crocuta crocuta*) (Busse, 1980; Cowlishaw, 1994; Jooste et al., 2013; Isbell et al., 2018).

Despite extensive work on baboon movement ecology and antipredator behaviour, few studies have tested the relative influence of resource-related environmental variables and predator activity on baboon space use within a multi-predator landscape. In this study, we took advantage of the largest camera trap-based monitoring project in Africa, the Snapshot Safari program (Swanson et al., 2015; Pardo et al., 2021). From a single study site in Tanzania, we analyzed 10,010 camera trap detections (baboon n = 1,531; lion n = 3,498; leopard n = 222; spotted hyena n = 4,759) to examine associations between environmental and predator covariates and olive baboon occupancy and detection probabilities. Specifically, we tested four hypotheses: Rugged terrain offers elevated sleeping refuges such as cliff faces and rocky outcrops that reduce nocturnal predation risk (Markham et al., 2015; Bidner et al., 2018). We therefore hypothesize that H1) baboon occupancy increases with terrain ruggedness index, a measure of topographic heterogeneity (Riley et al., 1999), at camera trap sites. Water availability strongly influences baboon spatial behaviour, with groups shifting their home ranges towards rivers and waterholes during dry periods (Johnson et al., 2015; Paietta et al., 2022; Hammond et al., 2025). We therefore hypothesize that H2) baboon occupancy will decrease as proximity to water sources decreases at camera trap sites. Denser vegetation structure provides greater sources of both food and refuge (Johnson et al., 2015; Markham et al., 2015; Bidner et al., 2018), including vertical escape routes in trees (Markham et al., 2015; Bidner et al., 2018; Suire et al., 2020). We therefore hypothesize that H3) baboon occupancy and detection probability will increase with vegetation greenness (NDVI, Normalized Difference Vegetation Index) at camera trap sites. Although lions and spotted hyenas only hunt and kill baboons opportunistically (Busse, 1980; Cowlishaw, 1994; Trinkel, 2010; Wentworth et al., 2011), as opposed to leopards, which actively hunt them (Busse, 1980; Cowlishaw, 1994; Trinkel, 2010; Wentworth et al., 2011; Isbell et al., 2018), predation remains the primary source of mortality in adult female and juvenile baboons (Cheney et al., 2004). We therefore hypothesize that H4) baboon occupancy will decrease as relative predator abundance (i.e., an index of predator activity at a camera-trap site inferred from the number of predator detections corrected for the deployment period of each camera trap) increases at camera trap sites.

## Methods

Reliable occupancy (ψ) and detection (*p*) probability estimates require a minimum of 35 camera traps for common species and over 150 for rare species (Kays et al., 2020), with deployments lasting a minimum of one month per camera. Accordingly, of the 32 study sites available from the large-scale, non-targeted camera-trap dataset from the ongoing Snapshot Safari monitoring program (Swanson et al., 2015), we selected a single study site, Serengeti National Park (SER; 2°19’S, 34°49’E; data publicly available at Dryad: http://dx.doi.org/10.5061/dryad.5pt92) in Tanzania (Figure 1), as its extensive camera trap coverage (n = 225) and long survey period from June 2010 to May 2013 (35 months), met the requirements for robust occupancy analysis.

**Figure 1.**
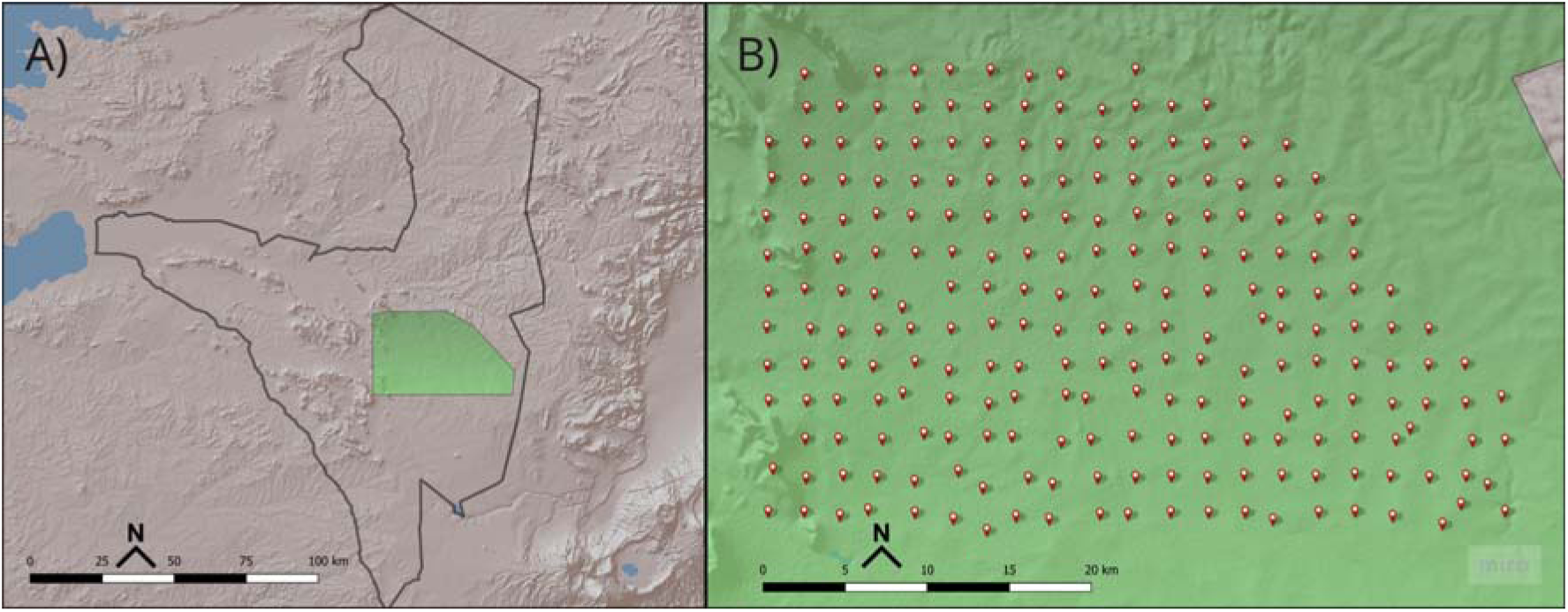
Study site (map adapted from Swanson et al (2015)): A) The Serengeti National Park boundary (black outline) located within Tanzania. The shaded green area represents the location of the study site; B) Camera trap locations (red dots, n = 219) within the boundaries of the study site.

SER spans approximately 25,000 km² within the East African savanna biome (Swanson et al., 2015), characterized by open grasslands interspersed with savanna woodlands (Anderson, 2008; Ogutu et al., 2008). Rainfall is bimodal, increasing from the southeast to the northwest, with the main wet season occurring between November and June and the dry season between July and October (Anderson, 2008; Ogutu et al., 2008; Mahony et al., 2021). Specifically, the short rains fall between November to December and the long rains between March and June (Anderson, 2008; Ogutu et al., 2008; Mahony et al., 2021).

### Camera trap data

A total of 225 Scoutguard (SG565) camera traps were deployed across an open subsection of SER following a systematic grid design, with each camera positioned at the center of a 5 km² cell (Swanson et al., 2015). Cameras were oriented according to local vegetation structure, avoiding tall grass to reduce the frequency of blank images (i.e. a frame or sequence of frames without animals Meek et al., 2014). Each camera was enclosed in a steel casing and mounted approximately 50 cm above the ground (Swanson et al., 2015), facing either north or south to minimize sun glare. Each camera was programmed to capture three images per motion trigger during the day and one at night (Swanson et al., 2015), with a medium sensitivity setting (50–75%) to limit false triggers from windblown vegetation. No lures were used to attract animals at any camera trap site.

Each detection event (i.e. a single image or consecutive image series initiated by an animal, regardless of the number of individuals Meek et al., 2014), was classified via Zooniverse (www.zooniverse.org) by trained volunteers of the general public. To ensure accuracy, each detection event was independently classified multiple times, using a plurality algorithm (i.e. final species identification was determined by voting consensus). The resulting dataset was then validated by experts, achieving a classification accuracy of 96.6% (Swanson et al., 2015).

### Statistical analyses

We derived environmental covariates and relative predator abundance metrics to test how they associated with baboon occupancy and detection probabilities.

We generated a baboon detection history matrix with the covariates: terrain ruggedness index, distance to the nearest river, seasonal NDVI values, and lion, leopard and spotted hyena relative predator abundance indices to evaluate their influence on baboon occupancy and detection probability. Camera trap data were aggregated into repeated seasonal sampling occasions for each camera trap site to allow estimation of detection probability, including a random intercept to account for repeated sampling of the same locations across seasons. Within each seasonal sampling period, site occupancy was assumed to remain constant. Detection probability was modelled as a function of NDVI only.

Terrain ruggedness index was calculated from a digital elevation model (DEM) with a 30.89 m by 30.92 m spatial resolution (Hijmans, 2023) using *QGIS* (QGIS.org, 2009) following the methods outlined by Riley et al (1999). The index measures elevation difference between a focal cell and its eight surrounding cells within a 3×3-pixel grid (Riley et al., 1999). For each camera trap site, we extracted the median index value within a 500 m radius buffer (0.78 km²) using the *terra* and *sf* packages in R (Pebesma & Bivand, 2018; Hijmans, 2023). This buffer size characterizes the immediate topographic conditions surrounding each camera trap site to avoid incorporating distant terrain features that may not affect local occupancy patterns.

As the study area lacks large permanent water bodies, distance to the nearest river was selected as a measure of water availability. Using the digital elevation model, we overlaid the surrounding river networks from the *HydroRivers* database (Lehner et al., 2013). Spatial vectors were processed in the *sf* package (Pebesma & Bivand, 2018) and converted to *SpatVector*s using the *terra* package (Hijmans, 2023) to calculate the distances (m) from each camera trap site to the nearest river.

We then estimated the vegetation productivity using the Normalized Difference Vegetation Index (NDVI), aggregated by 3-month seasonal periods, aligned with the camera trap deployment. The seasonal periods were defined as Wet Shoulder (December–February), Wet (March–May), Dry (June–August), and Dry Shoulder (September–November) based on the short and long rainfall patterns of the area (Anderson, 2008; Mahony et al., 2021; Ogutu et al., 2008). For each seasonal period, we calculated the median NDVI from each camera site from the MODIS MOD13A3 Version 6.1 dataset (Didan, 2021), which provides monthly NDVI data at a 500 m spatial resolution.

To examine how predator presence affects the occupancy and detection probability of olive baboons, we calculated the relative predator abundance as indices for each predator species (lions, leopards and spotted hyenas) (Rovero & Marshall, 2009). Relative predator abundance was calculated as an index of predator activity at a camera trap site, inferred from the number predator detections corrected for the deployment period of each camera trap. Separate models were run for each predator species to test species-specific influences on baboon occupancy due to the limited data and the potential collinearity between predators species (Périquet et al., 2015; Miller et al., 2018; Jones et al., 2021; Everatt et al., 2026)

To minimize pseudoreplication from multiple triggers of the same individual when calculating relative predator abundance of a species, we applied a 30-minute time-to-independence interval to all focal predator detections (lion, leopard and spotted hyena), retaining only the first detection within any 30-minute window (Linkie & Ridout, 2011; Searle et al., 2021). When a different species interrupted a predator sequence, subsequent events were considered independent detections (Swanson et al., 2015). This time to independence interval was not applied to our baboon detections as baboon presence in the area was not reliant on prey abundance.

Models were run using the R package *ubms* v.1.2.7 (Kellner, 2021) with default vague priors. Continuous covariates were standardised (mean = 0, SD = 1) to improve convergence and comparability. Each model was estimated with four MCMC chains with 6,000 iterations, discarding the first 3000 iterations per chain as burn-in. Convergence was assessed via the Gelman–Rubin statistic (□hat < 1.1) and visual inspection of trace plots. Model fit was evaluated using a MacKenzie-Bailey chi-square goodness-of-fit test (*p* > 0.05). A covariate was considered to have a strong predictive influence when its 95% credible intervals (CrI) did not overlap zero in either direction.

## Results

All three occupancy models (each including a different predator: lion, leopard, or spotted hyenas) converged successfully (RC = 1.00 for all parameters; effective sample size (n_eff) > 950 for all parameters). The models showed similar information criterion values (LOOIC: lion = 8,123.26, leopard = 8,120.31, spotted hyena = 8,123.26). However, goodness-of-fit tests indicated a poor overall fit (lion: MacKenzie-Bailey χ**²** = 3.08 × 1024, p = 0; leopard: MacKenzie-Bailey χ**²** = 2.63 × 1024, p = 0; hyena: MacKenzie-Bailey χ**²** = 4.02 × 1024, p = 0).

### Environmental covariate relationships with baboon occupancy

Baboon occupancy strongly increased with terrain ruggedness across all predator models (Figure 2). Effect sizes for environmental covariates were highly consistent across predator-specific models. For every one-SD increase in terrain ruggedness, the log-odds of occupancy increased by β = 1.46 in both the lion (95% CrI [0.80, 2.26]) and leopard models (95% CrI [0.80, 2.24]), and by β = 1.50 in the spotted hyena model (95% CrI [0.84, 2.30]). Conversely, baboon occupancy strongly decreased with increasing distance to rivers (DTR) across all predator models (Figure 2). For every one-SD increase in DTR, the log-odds of occupancy decreased by β = −1.14 in the lion model (95% CrI [-1.84, −0.50]), β = −1.13 in the leopard model (95% CrI [-1.79, −0.51]), and β = −1.15 in the spotted hyena model (95% CrI [-1.84, −0.52]).

**Figure 2.**
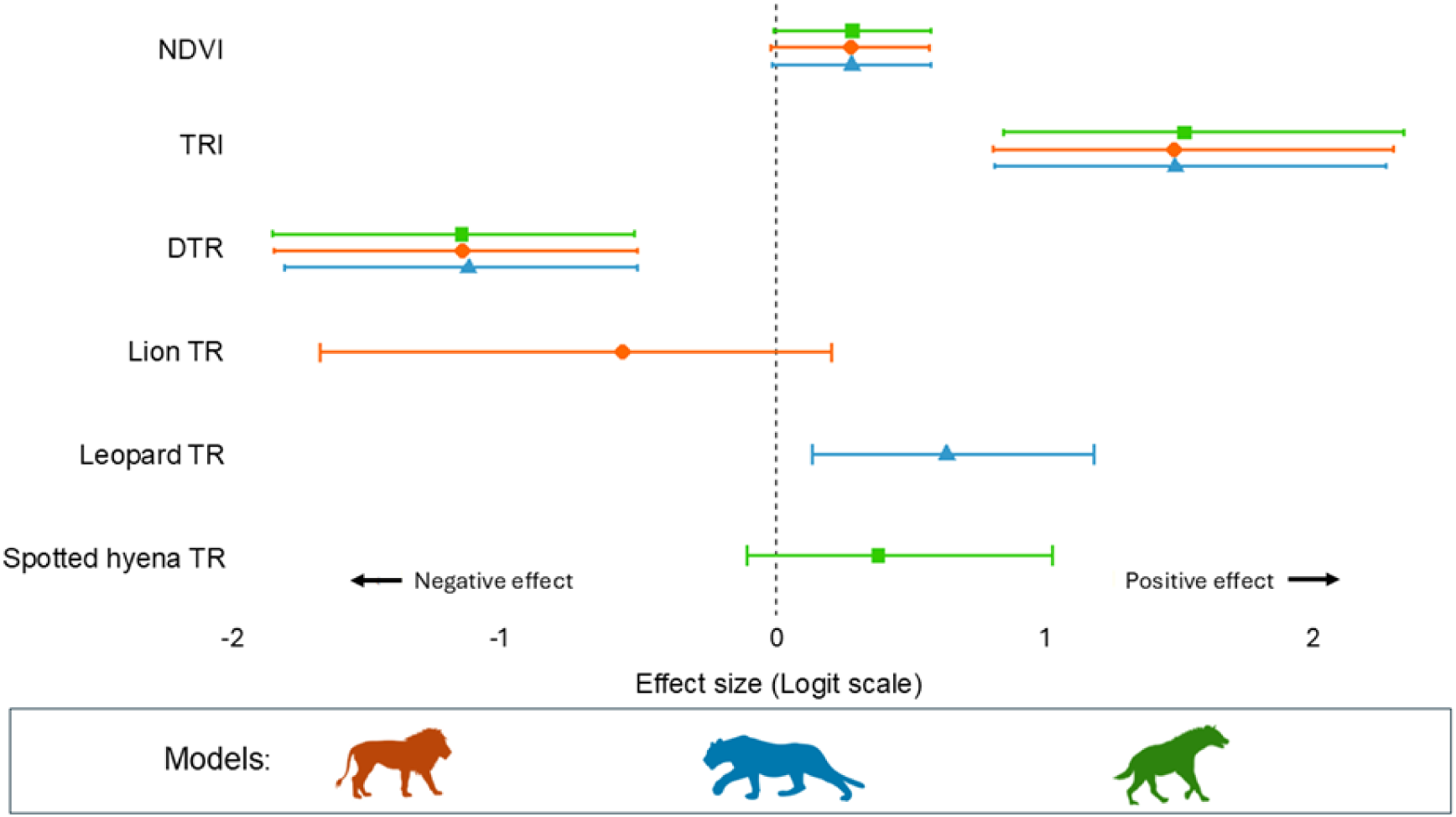
Posterior estimates of covariate influence on olive baboon occupancy from three bayesian occupancy models in the Serengeti ecosystem, Tanzania. Points represent posterior means with the horizontal bars indicating 95% credible intervals (CrI). The vertical dashed line at zero indicates no influence. Covariates with 95% CrI not overlapping zero are considered to have a strong (above zero) or weak (below zero) influence on occupancy.

Baboon occupancy showed a positive association with vegetation greenness (NDVI); however credible intervals overlapped zero across all three predator models (Figure 2). For every one-SD increase in NDVI, the log-odds of occupancy increased by β = 0.28 in the lion model (95% CrI [-0.01, 0.56]), β = 0.28 in the leopard model (95% CrI [-0.01, 0.57]), and β = 0.28 in the spotted hyena model (95% CrI [-0.01, 0.57]).

### Predator relative abundance relationships with baboon occupancy

Relative predator abundance showed contrasting associations with baboon occupancy (Figure 2). Baboon occupancy tended to decrease with lion trapping rate, though credible intervals overlapped zero (β = −0.56, 95% CrI [-1.67, 0.21]). Contrary to our predictions, baboon occupancy increased with leopard trapping rate, with credible intervals not overlapping zero (β = 0.63, 95% CrI [0.14, 1.17]). Finally, baboon occupancy showed a weak positive association with spotted hyena trapping rate, although credible intervals overlapped zero (β = 0.38, 95% CrI [-0.10, 1.01]).

### Environmental covariate relationships with baboon detection

NDVI showed a consistent positive association with baboon detection probability across all three predator models (Figure 3). For every one-SD increase in NDVI, the log-odds of detection increased by β = 0.14 in the lion model (95% CrI [0.07, 0.22]), β = 0.14 in the leopard model (95% CrI [0.07, 0.22]), and β = 0.14 in the spotted hyena model (95% CrI [0.07, 0.21]).

**Figure 3.**
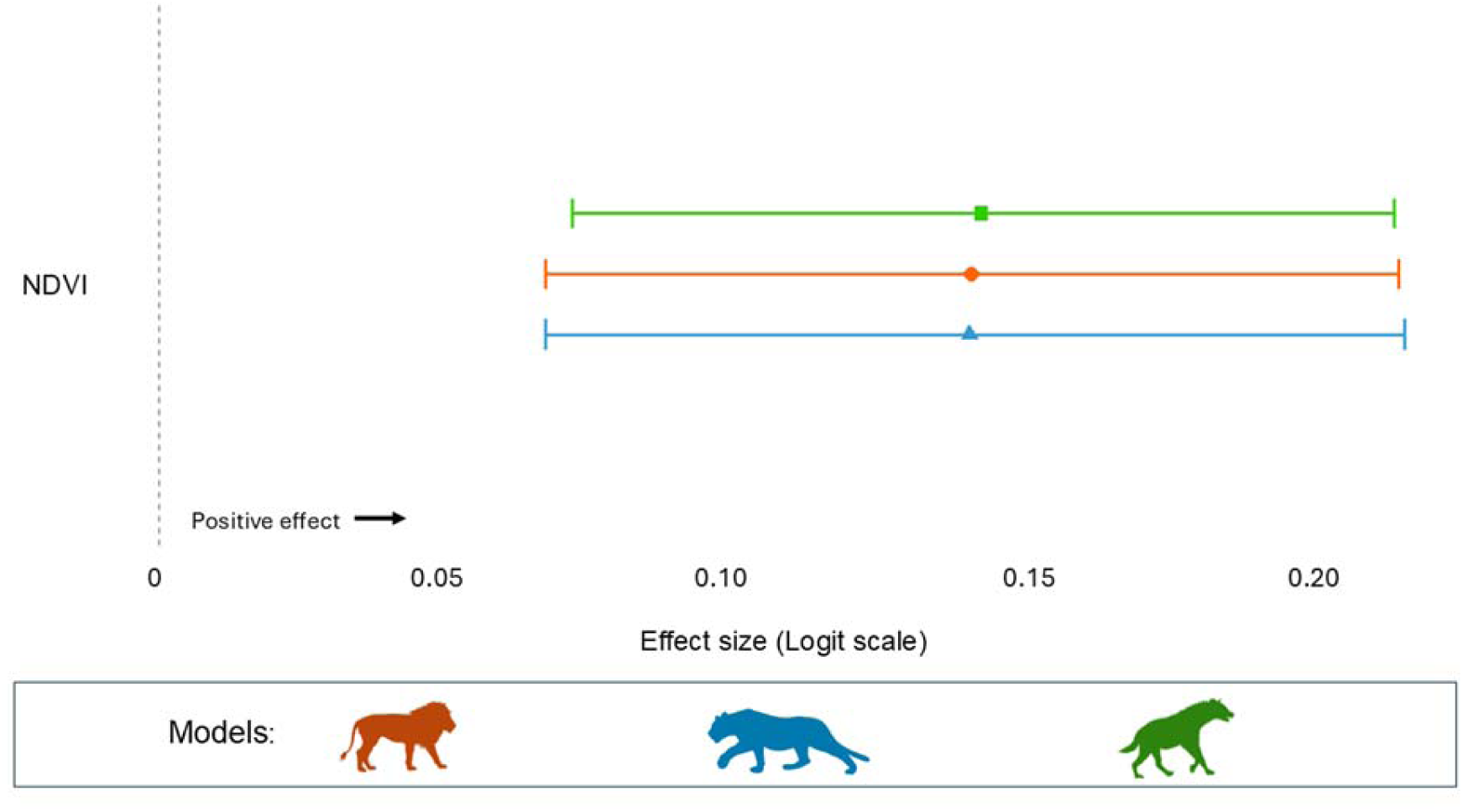
Posterior estimates of covariate influence on olive baboon detection from three bayesian models in the Serengeti ecosystem, Tanzania. Points represent posterior means with the horizontal bars indicating 95% credible intervals (CrI). The vertical dashed line at zero indicates no influence. Covariates with 95% CrI not overlapping zero are considered to have a strong (above zero) or weak (below zero) influence on detection.

## Discussion

Using camera trap data from a large-scale monitoring program in the Serengeti, Tanzania, we studied the influence of three environmental covariates (terrain ruggedness, distance to rivers, and vegetation greenness) and the relative predator abundance of three principal predators (lion, leopard, and spotted hyena) on olive baboon occupancy and detection probability. While previous studies have analysed baboon habitat selection in relation to environmental factors (Cowlishaw, 1997a; Johnson et al., 2015; Suire et al., 2021; Paietta et al., 2022), or predation effects separately (Cowlishaw, 1997b; Markham et al., 2015; Isbell et al., 2018; Allan et al., 2024), this study explicitly integrated both sets of drivers within occupancy models to assess their relative influence on olive baboon spatial distribution. Across models, baboon occupancy strongly increased with terrain ruggedness and strongly decreased with an increase in distance to rivers. While NDVI showed only marginal associations with baboon occupancy (Figure 2), we detected a stronger influence on baboon detection probability (Figure 3). Relative predator abundance showed varied influences on baboon occupancy between models: lion and spotted hyena relative abundance were not strongly associated with baboon occupancy; in contrast, baboon occupancy strongly increased with relative leopard abundance.

### Environmental correlates of baboon occupancy

The strong associations between baboon occupancy and environmental covariates highlight the importance of landscape structure and resource distribution in olive baboon habitat use (Altman et al., 1970; Bettridge et al., 2010; Johnson et al., 2015; Strandburg-Peshkin et al., 2017; Suire et al., 2021; Paietta et al., 2022). As a diurnal primate, the increase in baboon occupancy with terrain ruggedness aligns with well-documented baboon behaviour, where steep rocky areas such as cliffs and ledges provide secure sleeping refuges that reduce vulnerability to nocturnal predation (Cowlishaw, 1997b; Schreier & Swedell, 2012; Markham et al., 2015; Bidner et al., 2018; Isbell et al., 2018; Suire et al., 2021). This terrain-dependent refuge strategy is widespread across taxa, having been reported in both other Old World monkeys, such as geladas (*Theropithecus gelada*) and in some ungulates such as klipspringers (*Oreotragus oreotragus*) and mule deer (*Odocoileus hemionus*), all of which rely on rugged topography as an effective antipredator response (Crook & Aldrich-Blake, 1968; Lingle, 2002; Druce et al., 2009).

Similarly, the strong decrease in baboon occupancy with an increase in distance to water reflects baboons’ dependence on surface water, particularly during dry seasons when groups shift their home ranges and sleeping sites toward water sources, which also provide access to riparian fruit trees and other herbaceous foods (Barton et al., 1992; Isbell et al., 2018; Paietta et al., 2022; Hammond et al., 2025). Reliance on riparian habitats is a common ecological driver across African mammals. Large herbivores such as elephants (*Loxodonta africana*) and Cape buffalo (*Syncerus caffer*) also show occupancy patterns strongly predicted by proximity to water (Sadie et al., 2012; Davis et al., 2023). However, because predator occurrence is similarly structured (Schuette et al., 2013), these areas often become zones of elevated predation risk (Sirot et al., 2016; Courbin et al., 2018; Paietta et al., 2022). Prey species—including baboons—must therefore balance the need to access essential resources with the increased likelihood of encountering predators. For baboons, this is managed through behavioural and coalitionary anti-predator strategies, such as sentinel behaviour, alarm calls, coordinated mobbing, and cohesion (Karpanty & Wright, 2007; Cowlishaw, 1997a; Gursky, 2007; Bezerra et al., 2008; Crofoot, 2013; Bidner, 2018).

The association with vegetation greenness (NDVI) and baboon occupancy was consistent across all three predator models (β ≈ *0.28*), however the seasonal fluctuations in rainfall may help to explain the credible intervals received (Fabricante et al., 2009). The observed association between occupancy and NDVI suggests that vegetation productivity may be related to olive baboon occupancy, potentially through increased food availability, greater vegetative cover for predator avoidance, and enhanced access to arboreal refuge strata (Bidner et al., 2018; Johnson et al., 2015; Markham et al., 2015). It is also highly probable that baboon occupancy at camera trap sites fluctuates seasonally, being stronger at camera trap sites with higher-NDVI values during drier seasons when vegetation greenness is less uniform across the study site (Johnson et al., 2015; Hammond et al., 2025). However, the uncertainty indicates that NDVI alone may not fully capture relevant resource heterogeneity, likely reflecting the coarse spatial resolution of greenness indices and the importance of fine-scale habitat features, as shown in earlier studies showing scale-dependent effects of vegetation productivity on primate ranging behaviour (Fehlmann et al., 2017; Suire et al., 2021).

### Leopard co-occurrence and shared habitat selection

The strong association between leopard abundance and baboon occupancy likely reflects shared habitat preferences rather than predator tolerance. Both species favour rugged terrain (Cowlishaw, 1997b; Schreier & Swedell, 2012; Miller et al., 2018; Suire et al., 2021) and densely vegetated areas (Balme et al., 2007; Isbell et al., 2018; Miller et al., 2018). Leopards rely on the provided cover to help facilitate ambush hunting and act as sleeping sites (Balme et al., 2007; Miller et al., 2018), while baboons use vertical structures for refuge from predators and increased visibility (Cowlishaw, 1997b; Schreier & Swedell, 2012; Markham et al., 2015; Bidner et al., 2018; Suire et al., 2021).

Despite leopards occasionally preying on baboon (Busse, 1980; Bidner et al., 2018; Isbell et al., 2018), successful kills are typically limited to vulnerable individuals or small groups (Suire et al., 2023; Jooste et al., 2013). Baboons are not preferred leopard prey (Hayward et al., 2006), largely due to their aggressive coalitionary responses, including male-driven defence, collective vigilance, and mobbing (Bettridge et al., 2010; Jooste et al., 2013; Isbell et al., 2018; Suire et al., 2023). Similar patterns of habitat overlap have been described in other African primates. Patas monkeys (*Erythrocebus patas*) avoid high-risk leopard habitats but overlap home ranges with lion ranges, demonstrating that co-occurrence does not necessarily equate to elevated predation exposure (Burnham & Riordan, 2013). Likewise, vervet monkeys (*Chlorocebus pygerythrus*) trade off access to food-rich riparian zones against heightened leopard risk (Willems & Hill, 2009). Together, these examples support the interpretation that primates may occupy high-risk habitats when the benefits of resource access and refuge availability outweigh the costs of predation risk (Sih, 2005; Riginos, 2015).

### Lions and spotted hyenas: habitat partitioning and weak associations

In contrast to our leopard findings, lion and spotted hyena showed no associations with olive baboon occupancy as their credible intervals both cross 0 (Figure 2). Both predator species prefer open to semi-open habitats with high densities of large (190–550 kg) to medium (56–182 kg) ungulate prey (Trinkel, 2010; Miller et al., 2018; Jones et al., 2021). Lions require some degree of concealment for successful hunting, with hunting success increasing in taller grass while avoiding denser woodland, as this reduces prey encounter rates and may hinder cooperative hunting (Périquet et al., 2015; Miller et al., 2018). As cursorial predators spotted hyenas are described as habitat generalists that occupy any habitat where prey are most abundant, without requiring specific vegetation cover (Trinkel, 2010; Wentworth et al., 2011; Périquet et al., 2015). The absence of strong associations likely reflects habitat partitioning as described by MacArthur (1958), where baboons may occupy structurally complex habitats that have limited overlap with the open landscapes favoured by lions and spotted hyenas (Périquet et al., 2015; Jones et al., 2021). When encounters do occur, baboons likely rely on the behavioural defenses described previously rather than complete spatial avoidance. Such strategies may be energetically more efficient than large-scale avoidance, particularly in multi-predator systems where complete spatial avoidance of all predators is not possible (Sih, 2005; Bettridge et al., 2010; Riginos et al., 2015). Moreover, weak spatial associations could arise if baboons mitigate risk through temporal rather than spatial partitioning, adjusting activity timing to avoid peak predator activity as observed in other diurnal primates (Gaynor et al., 2019).

### Detection probability patterns

Detection probability increased with NDVI across all predator models. This suggests that olive baboons were more frequently detected in areas with higher vegetation density, likely reflecting increased ground-level foraging activity when ground food resources are abundant (Johnson et al., 2015; Hammond et al., 2025). Increased ground activity would therefore increase the probability of camera detections. Similar, camera-trap studies on both hamadryas and yellow baboons (*Papio cynocephalus*), as well as other primate taxa, such as vervet monkeys have shown higher detection rates in areas with greater vegetation greenness or canopy complexity, reflecting both resource concentration and improved thermal buffering (Pochron, 2000; Fehlmann et al., 2017; Paietta et al., 2022).

### Potential limitations

Several limitations must be acknowledged that may have constrained our ability to fully detect baboon-predator relationships. First, as the study was initially designed to target lions in the Serengeti (Swanson et al., 2015), the spatial scale of the camera trap placement (5 × 5 km grid) may have been too coarse to detect fine-scale movement patterns of baboons. Increasing camera trap numbers may therefore be necessary to improve detection rates of both baboons and predators (Si et al., 2014). It has been shown that adding a second camera trap at each camera trap site can increase the detection probability and precision of occupancy estimates by as much as 400%, when targeting predators such as coyotes (*Canis latrans*) and the Eurasian lynx (*Lynx lynx*) (Evans et al. 2019; Hofmeester et al., 2021).

Second, detection methodology may have limited our ability to accurately capture baboon occupancy. Ground-based camera traps and line transect surveys are effective at detecting terrestrial mammals, however they are less effective at capturing the true activity levels or space use of semi-arboreal species such as baboons (Bowler et al., 2016; Moore et al., 2020). Moore et al. (2020) found that by combining ground and arboreal camera traps they detected 91% of the targeted aboral and semi arboreal mammal species, compared to 71% for ground cameras alone, showing highest detection probabilities under different methods. Similarly, Bowler et al. (2016) demonstrated that arboreal cameras detected cryptic nocturnal species missed by ground-based methods. Therefore, integrating both arboreal cameras or GPS collars of known groups could refine our understanding of daily movement patterns and validate whether ground-based detectability reflects actual habitat use (Hopkins & Milton, 2016; Isbell et al., 2018), thereby improving detection rates and spatial representation of baboons.

Third, despite adequate model convergence and draw estimates (n_eff > 950), goodness-of-fit tests indicated a poor overall fit across all three predator models (MacKenzie-Bailey χ² > 10²□, p = 0 for all models). This suggests potential unmodelled heterogeneity in baboon space use, indicating that while our models capture the relative importance of key predictors, they may not fully explain absolute occupancy patterns (MacKenzie & Bailey, 2004; Kays et al., 2021). However, as the aim of our study was not to predict baboon occupancy but rather investigate the associations with baboon occurrence, we are still able to make ecological inferences regarding the effects of the selected covariates. (MacKenzie & Bailey, 2004; Kays et al., 2021; Tredennick et al., 2021; Stewart et al., 2022).

## Conclusion

This study helps contribute to broadening our understanding of predator-prey interactions and the ecological drivers that shape primate occupancy patterns in multi-predator landscapes. Our findings demonstrate that resource-related environmental features—particularly access to rugged terrain and water sources—are primary and consistent influences of olive baboon spatial distribution in the Serengeti, maintaining their strength regardless of predator presence. This suggests that for a behaviourally flexible primate species with diverse antipredator strategies, the priority to access essential resources may in most cases surpass the proactive spatial avoidance of predators. The positive association between leopard and baboon occupancy likely reflects habitat co-occurrence driven by shared habitat preferences rather than species interactions, while the absence of relationships with lion and spotted hyena suggests habitat partitioning or reliance on behavioural defences rather than large-scale spatial avoidance. These patterns help to outline that in multi-predator system where complete spatial avoidance of all predators is impossible, baboons may prioritize access to critical resources while employing alternative antipredator strategies when encounters occur. However, the poor model fit across our three predator models suggests that future research incorporating additional methods of data collection would provide a more comprehensive insight into the drivers of baboon habitat selection.

## Authors’ contributions

Conceptualization: VR, FP, TH – Methodology: NVR, TH, VR, FP – Formal analysis: NVR, TH, VR, FP – Investigation: NVR, VR, FP – Writing: Original draft: NVR, VR, FP, TH – Writing: Review & Editing: NVR, TH, VR, FP – Supervision: TH, VR, FP – Funding acquisition: VR

## Acknowledgments

The authors thank the CNRS for funding this project. We thank the Snapshot Serengeti project for providing access to the camera-trap data used in this study, and in particular Craig Packer and the Snapshot Serengeti team for their long-term efforts in data collection and curation. We are also grateful to the many volunteers who contributed to data classification through the Snapshot Serengeti platform. We thank Serengeti National Park authorities for permitting research in the park.

## Inclusion and diversity statement

The authors are committed to fostering an environment of open dialogue, respect, and cultural inclusion. Our collaboration between African and European researchers reflects our dedication to international scientific exchange and diverse perspectives. Within our research team and laboratory, we actively promote a supportive and inclusive environment. Our first author lives with temporal lobe epilepsy, a condition that affects both memory and language. We acknowledge the unique challenges this presents in research and take meaningful steps to ensure that our team remains understanding and accommodating to this disability. We firmly oppose all forms of discrimination based on sex, gender, race, or any other identity marker. Our partnership is grounded in fairness and merit, and we evaluate all contributions based on the quality and impact of the work, not on the individual’s background or identity.

## Data availability statement

Pictures from the Snapshot Serengeti program are available from the Labelled Image Library of Alexandria – Biology and Conservation: http://lila.science/datasets/snapshot-serengeti. All classification data and metadata are publicly available at Dryad: http://dx.doi.org/10.5061/dryad.5pt92.

## Code availability statement

The code to analyze and reproduce this study has been deposited in Zenodo and is available online at https://doi.org/10.5281/zenodo.18457909

### Conflict of interest

The authors declare that they have no conflict of interest.

### Ethical approval

Ethics approval was not required for this study according to local legislation [Parks and Wildlife Act].

Consent to participate: Not applicable.

Consent for publication: Not applicable.

## Notes

http://dx.doi.org/10.5061/dryad.5pt92

### Competing Interest Statement

The authors have declared no competing interest.

